# Muscle coordination retraining inspired by musculoskeletal simulations: a study on reducing knee loading

**DOI:** 10.1101/2020.12.30.424841

**Authors:** Scott D Uhlrich, Rachel W Jackson, Ajay Seth, Julie A Kolesar, Scott L Delp

## Abstract

Humans typically coordinate their muscles to meet movement objectives like minimizing energy expenditure. In the presence of pathology, new objectives gain importance, like reducing loading in an osteoarthritic joint, but people often do not change their muscle coordination patterns to meet these new objectives. Here we use musculoskeletal simulations to identify simple changes in coordination that can be taught by providing feedback of electromyographic activity to achieve a therapeutic goal—reducing joint loading. Our simulations predicted that changing the relative activation of the redundant ankle plantarflexors could reduce knee contact force during walking, but it was unclear whether humans could re-coordinate redundant muscles during a complex task like walking. With simple biofeedback of electromyographic activity, healthy individuals reduced the ratio of gastrocnemius to soleus muscle activation by 25±15% (p=0.004). The resulting “gastrocnemius avoidance” gait pattern reduced the late-stance peak of simulation-estimated knee contact force by 12±12% (p=0.029). Simulation-informed muscle coordination retraining could be a promising treatment for knee osteoarthritis and a powerful tool for optimizing coordination for a variety of rehabilitation and performance applications.

## Introduction

The human musculoskeletal system is equipped with more than the minimum number of muscles needed to produce movement. This muscular redundancy allows the central nervous system to optimize movement to meet task-specific performance goals, such as walking efficiently, climbing safely, or running quickly. A vast number of muscle coordination strategies can generate the same motion, but each strategy results in different internal quantities such as joint loading, metabolic cost, or tendon strain. During walking, for example, healthy humans are thought to select a coordination strategy that optimizes performance metrics such as reduced metabolic cost^1,2^ and increased stability^3^. In the presence of pathologies such as osteoarthritis, stroke, or ligament injury, new performance metrics may become increasingly important. However, humans do not always adopt new coordination strategies that optimize for these new metrics, potentially due to the lack of robust and timely feedback mechanisms or insufficient exploration of new coordination patterns^4^. Musculoskeletal simulations allow us to explore the relationships between neuromuscular control, kinematics, and clinically-relevant metrics like joint loading to identify more favorable coordination strategies that would be challenging for humans to discover without guidance. This study examines the utility of simulation-guided muscle coordination retraining in designing a joint-offloading intervention for individuals with knee osteoarthritis.

Reducing compressive loading in the knee is a target for many non-surgical treatments for knee osteoarthritis, due to the relationship between excessive loading and osteoarthritis symptoms^5^ and progression^6–8^. During walking, compressive knee contact force (KCF) reaches between 2-4 times bodyweight (BW; Figure 1). Fifty to seventy-five percent of this force is in reaction to tensile muscle forces across the joint^9–12^, and the remaining force is the resultant force from inverse dynamics, called the intersegmental force. Interventions like osteotomy^13^, bracing^14^, and kinematic gait retraining^15–17^ aim to reduce knee loading by altering the intersegmental force. However, these interventions do not target, and often increase, the large muscle contribution to KCF^18^. Additionally, despite the relationship between knee loading and pain, individuals with osteoarthritis do not naturally select coordination strategies that minimize joint loading^19–21^. Even when provided with real-time feedback of KCF, individuals are not able to alter their muscle coordination to reduce KCF^22^. This suggests that the complex dynamics relating motor control to KCF, not exclusively a lack of feedback, could limit the ability of individuals with osteoarthritis to adopt a load-reducing coordination strategy.

**Figure 1:**
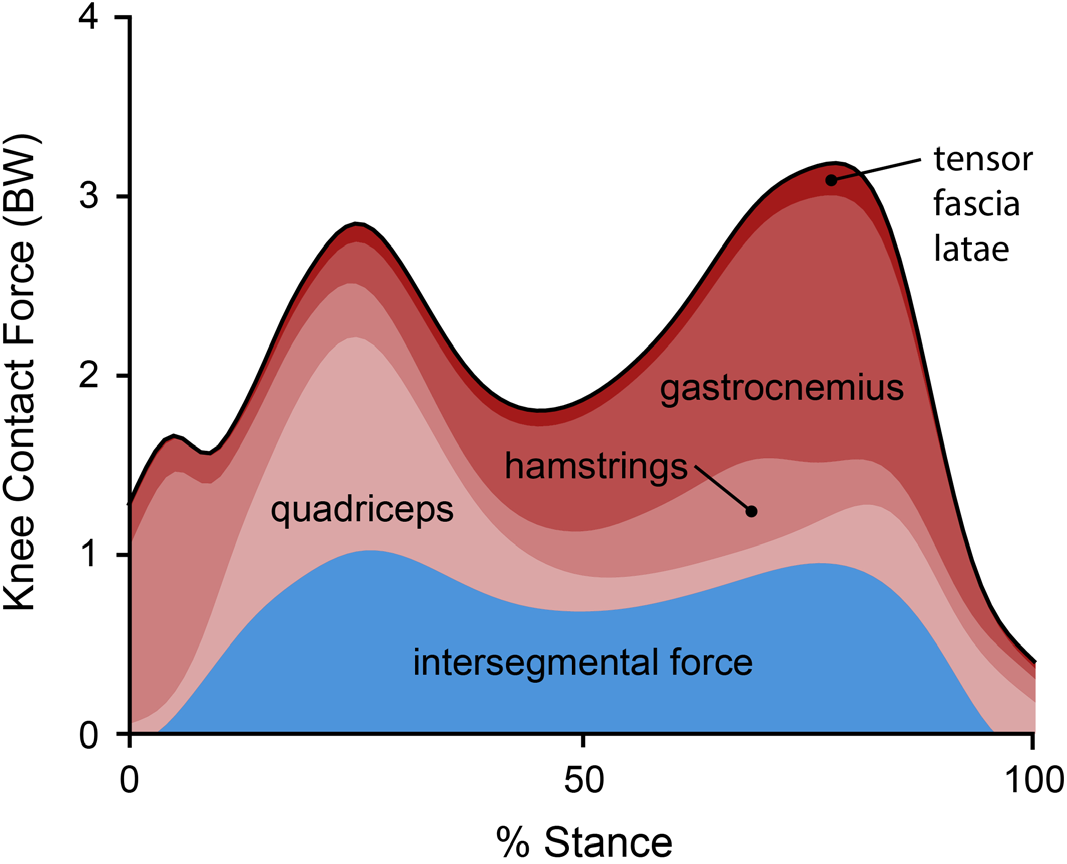
The relative contribution of the intersegmental reaction force (resultant force from inverse dynamics) and muscle forces to knee contact force. Muscle force contributions are dominated by the hamstrings during the first 10% of stance, the quadriceps from 10-40% stance, and the gastrocnemius from 40-90% stance. The components of knee contact force are shown for the 10 healthy subjects in this study with muscle activations estimated by minimizing the sum of squared activations using static optimization.

Musculoskeletal simulations can identify coordination strategies that reduce KCF. The second, or late-stance, peak of knee contact force (KCF_P2_) is sensitive to changes in muscle coordination and can, in theory, be altered by several times bodyweight without changing kinematics^20,23,24^. A coordination strategy that minimizes KCF_P2_ can be achieved by altering the activation of every lower-extremity muscle^20,24^, but this solution is too complicated to learn. Solutions that involve simpler changes in muscle coordination, like reducing the activation of the gastrocnemius^11,20,23–25^, are likely easier to learn and can still substantially reduce KCF_P2_. While simulations suggest that a “gastrocnemius avoidance” coordination strategy could effectively reduce KCF_P2_, it remains unclear which redundant muscles would need to compensate for the reduction in gastrocnemius activation. Furthermore, it is unknown whether humans can learn to change the relative activation of redundant muscles during a complex task like walking.

Electromyography (EMG) biofeedback is an effective tool for exploring the limits of volitional motor control. During walking, biofeedback of the tibialis anterior or gastrocnemius can help restore normal ankle dorsiflexion or plantarflexion moments for individuals with stroke or cerebral palsy^26,27^. Humans can also gain selective control over several motor units within a single biofeedback session^28,29^, but most individuals lose this selective control during dynamic tasks^30^. EMG biofeedback can also aid in selecting kinematic strategies that change the relative activation of redundant muscles during physical-therapy exercises^31,32^. There has been limited work, however, exploring whether humans can change the relative activation of redundant muscles without changing kinematics during dynamic tasks like walking.

The purpose of this study was to design and implement a muscle coordination retraining intervention that teaches individuals to reduce knee contact force. We first used musculoskeletal simulations to identify a subset of muscles to target with biofeedback that would reduce knee contact force without changing joint kinetics (Figure 2a). Based on the simulation results, we designed an EMG biofeedback intervention to test whether healthy individuals could learn to change the coordination of redundant muscles during walking (Figure 2b). We then used EMG-informed simulations to evaluate whether participants reduced their knee contact force when walking with the new coordination pattern (Figure 2c). More generally, we describe a framework for using simulations to design simple biofeedback interventions that teach individuals new coordination patterns that have a therapeutic or performance-enhancing effect.

**Figure 2:**
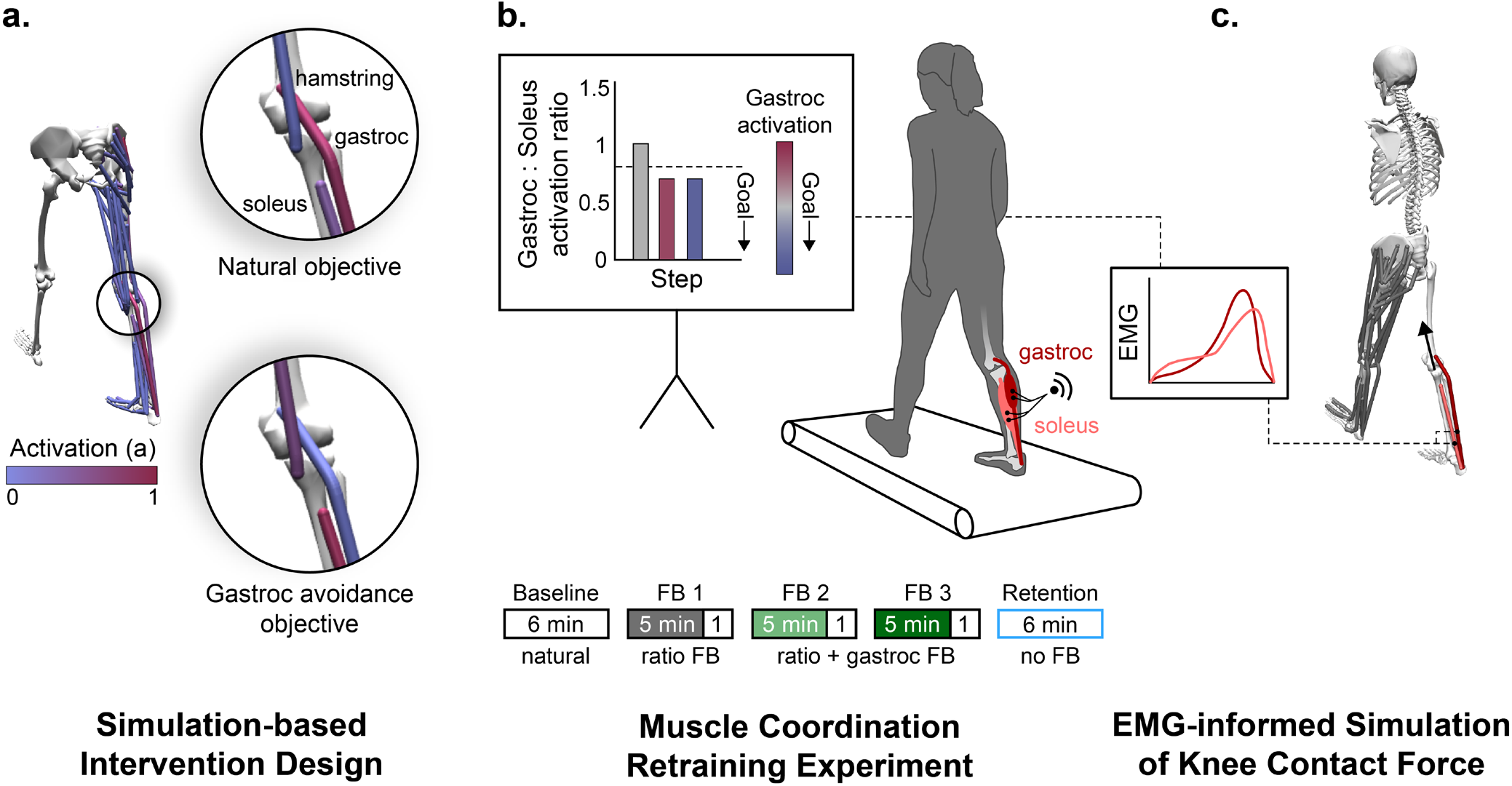
**(a)** To design the biofeedback intervention for teaching individuals to walk without activating their gastrocnemius (gastroc), we simulated a healthy adult walking using both a natural and a gastrocnemius avoidance objective function (Equations 1 and 2) in the static optimization muscle redundancy solver. **(b)** Based on the simulation results, we provided individuals with electromyography (EMG) biofeedback that instructed them to change the coordination of their ankle plantarflexor muscles. Participants performed five walking trials: a baseline trial (natural walking), three trials with visual biofeedback, and a retention trial without feedback. During all three feedback (FB) trials, visual biofeedback instructed participants to reduce their gastrocnemius-to-soleus activation ratio (bar magnitude). During the final two feedback sessions, additional feedback was provided, instructing participants to also reduce their average gastrocnemius activity (bar color). We analyzed steps from the final minute of each trial, during which, no feedback was provided. **(c)** We generated EMG-informed static optimization simulations to assess the effect of a gastrocnemius avoidance coordination pattern on knee contact force. The simulated gastrocnemius-to-soleus activation ratio was constrained to match the ratio measured with EMG.

## Results

### Simulation-based intervention design

To design the coordination retraining intervention, we sought to understand the compensatory muscle activations that are necessary to generate normal walking kinetics with minimal gastrocnemius activation (Figure 2a). We simulated healthy walking kinetics with two different static optimization objective functions: a natural coordination objective that is a surrogate for metabolic energy^33,34^ (Equation 1) and a gastrocnemius avoidance objective that additionally penalizes gastrocnemius activation^20^ (Equation 2).

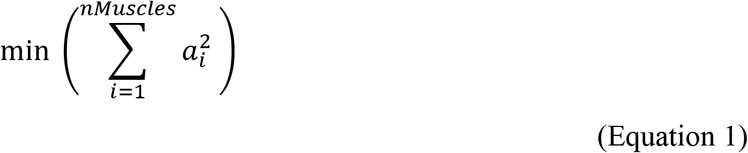

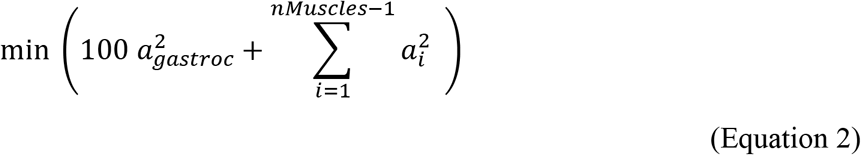

These simulations showed that walking without activating the gastrocnemius requires increased force generation from the soleus, hamstrings, and hip flexors (Figure 3). Increased soleus force supplements the ankle plantarflexion moment generated by gastrocnemius, but unlike the gastrocnemius, the soleus does not cross the knee, allowing it to plantarflex the ankle without compressing the knee. Increased hamstrings force supplements the knee flexion moment generated by the gastrocnemius, but the hamstrings have, on average, a knee flexion moment arm that is 1.7 times greater than the gastrocnemius^35^, allowing these muscles to generate the same moment with less force. The large changes in the force required by the ankle plantarflexors coupled with their essential role in generating normal walking kinematics^36,37^ led us to focus the biofeedback intervention on the relative activation of the gastrocnemius and soleus muscles. With the goal of teaching participants to reduce gastrocnemius force without reducing their plantarflexion moment, we designed biofeedback that instructed participants to reduce gastrocnemius activation and increase soleus activation.

**Figure 3:**
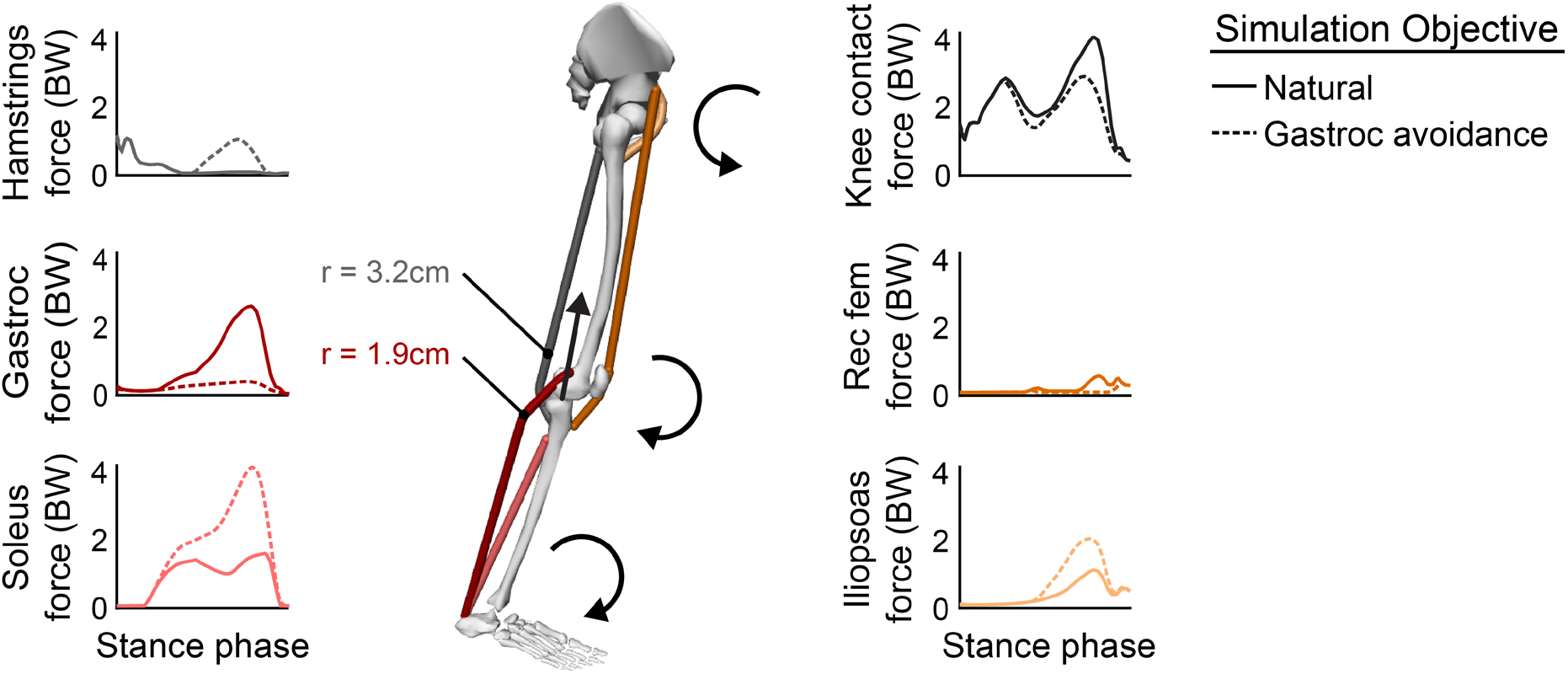
Simulations of identical joint kinetics (n=1) with a natural (Equation 1) and a gastrocnemius avoidance (Equation 2) static optimization objective function. During late stance, the muscles are generating ankle plantarflexion, knee flexion, and hip flexion moments. The gastrocnemius avoidance coordination pattern requires the soleus and hamstrings muscles to generate more force to compensate for the ankle plantarflexion and knee flexion moment-generating functions of the gastrocnemius. Due to their greater knee flexion moment arms (r), the hamstrings can generate the knee flexion moment with less force than the gastrocnemius, thereby reducing the muscle contribution to knee contact force. This increased hamstring force generates an antagonistic hip extension moment which is counteracted by an increase in iliopsoas force. Muscle-group-averaged moment arms were computed as a weighted average of the moment arms (hamstrings: biceps femoris long head and short head, semitendinosus, semimembranosus; gastrocnemius: medial and lateral heads) at 75% of the stance phase, weighted by each muscle’s optimal force in the musculoskeletal model^35^.

### Muscle coordination retraining experiment

We conducted an experiment to explore whether humans could learn to change the relative activation of their gastrocnemius and soleus muscles during walking when given visual biofeedback. During a single session, 10 healthy adults performed five six-minute walking trials on an instrumented treadmill: a baseline trial, three feedback trials, and a retention trial (Figure 2b). During all feedback trials, a real-time bar plot instructed participants to reduce their gastrocnemius-to-soleus activation ratio (Equation 3):

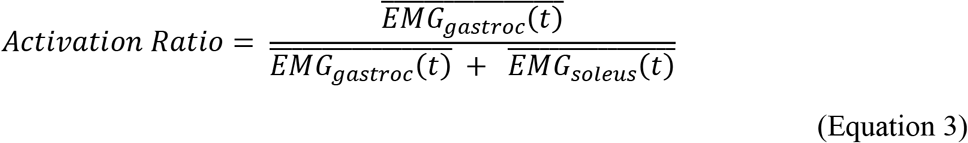

where 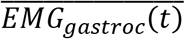 and 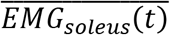 are the average EMG linear envelopes over the stance phase for the medial gastrocnemius and soleus, respectively. During the final two feedback trials, participants were also given feedback to reduce their average gastrocnemius activation to prevent them from reducing the activation ratio by only increasing soleus activation. During the retention trial, participants were instructed to retain their learned coordination pattern but were not given feedback.

Following the first trial, where participants were only given feedback of the activation ratio, they reduced their activation ratio by 22±12% (p<0.001), but did not significantly reduce gastrocnemius activation (Figure 4a). By adding gastrocnemius activation feedback during the second and third feedback trials, we sought to teach participants to reduce their activation ratio by reducing gastrocnemius activation. Following the third feedback trial, participants reduced gastrocnemius activation by 17±19% (p=0.033) compared to baseline. Finally, we investigated whether individuals could retain their new coordination pattern after six minutes of walking without feedback. At the end of the retention trial, participants retained a 25±15% (p=0.004) reduction in activation ratio and a 17±21% (p=0.033) reduction in gastrocnemius activation. Despite this change in activation ratio, the average ankle moment only changed by 3±14% (Figure 4e) during the retention trial compared to baseline, which trended towards statistical equivalence within one baseline standard deviation (p=0.063).

**Figure 4:**
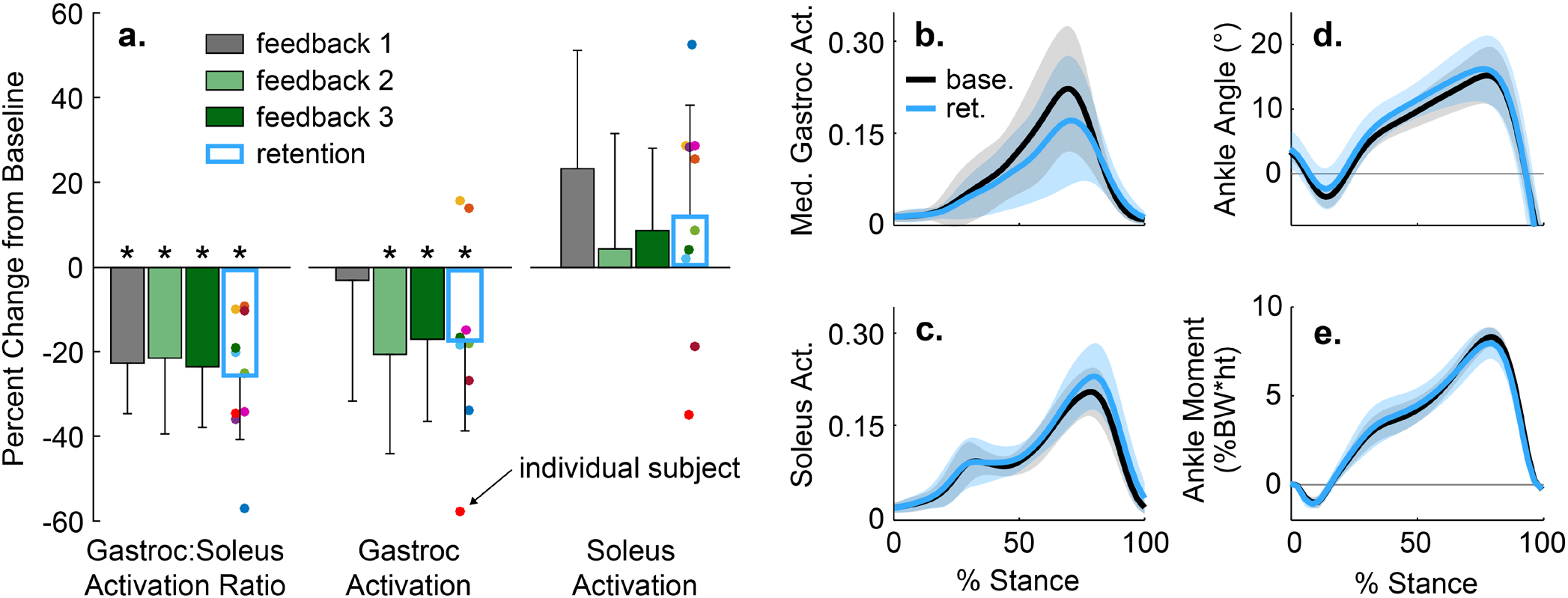
**(a)** The average (bar), and standard deviation (error bar) of changes in muscle activation measured with electromyography (n=10). Participants reduced their gastrocnemius (gastroc) to soleus activation ratio by 22±12% (p<0.001) following the initial feedback session and retained a 25±15% (p=0.003) reduction at the end of the retention trial. Participants reduced average gastrocnemius activation by 17±19% (p=0.033) following the third feedback session and retained a 17±21% (p=0.033) reduction. **(b-e)** The average (line) and standard deviation (shading) of medial gastrocnemius and soleus muscle activity and ankle mechanics for the baseline (base.) and retention (ret.) trials. Despite the 25±15% change in activation ratio from the baseline to retention trial, the stance-phase-averaged ankle moment only changed by 3±14%, which trended towards being equivalent to baseline within one baseline standard deviation (p=0.063).

### EMG-informed simulation of knee contact force

We evaluated whether the gastrocnemius avoidance coordination pattern that participants adopted achieved the desired therapeutic objective: a reduction in simulation-estimated KCF_P2_. Using static optimization, we simulated five gait cycles from the final minute of both the baseline and retention trials. To capture changes in ankle plantarflexor muscle activity between conditions, we constrained the gastrocnemius-to-soleus activation ratio at each time step in the simulation to match the corresponding ratio measured with EMG (Figure 2c). The simulated activations qualitatively matched EMG patterns for the major muscle groups crossing the knee and ankle (Figure 5).

**Figure 5:**
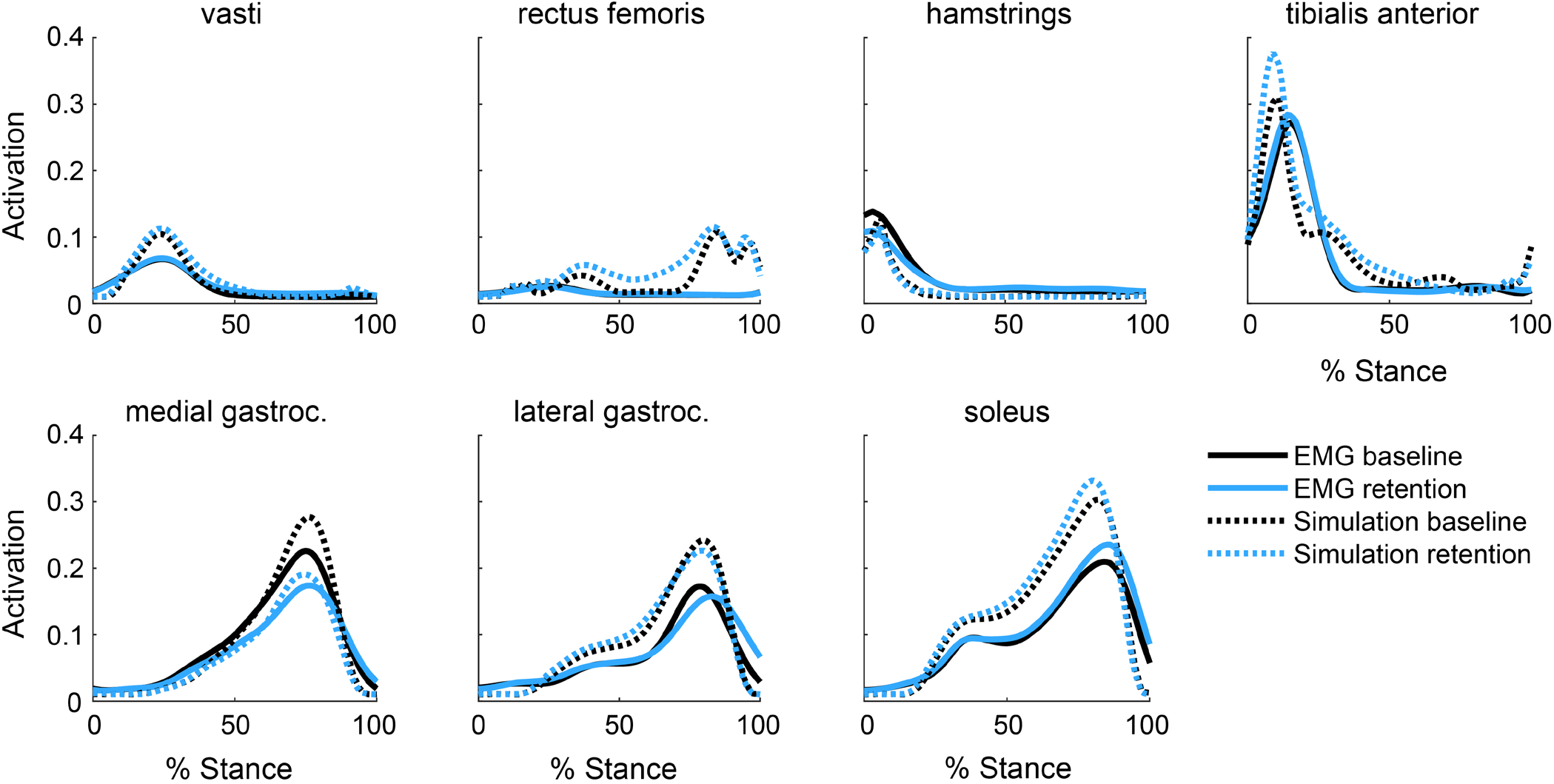
Electromyography (EMG) linear envelopes with a 40ms electromechanical delay compared to simulated muscle activations for the baseline and retention trials for all participants (n=10). Despite magnitude differences between EMG and simulated activations of the gastrocnemius and soleus, the relative changes between trials are consistent, indicating that our EMG-informed static optimization technique captured the changes in ankle plantarflexor muscle activity measured with EMG. The activations of the muscles in the top row were not informed by EMG in the simulation. The shape, timing, and between-trial changes in simulated activation matched EMG for these muscles, with the exception of the rectus femoris.

To determine the efficacy of the intervention, we analyzed joint contact forces for the eight individuals who retained a reduction in late-stance gastrocnemius activation during the retention trial. On average, these individuals reduced their KCF_P2_ by 0.38±0.39 BW (12±12%, p=0.029) during the retention trial compared to baseline (Figure 6a). The 0.25±0.30 BW reduction in the gastrocnemius contribution to KCF was the main contributor to reduced KCF_P2_ (Figure 6b). Contrary to the prediction by the intervention-design simulation (Figure 3), neither the hamstrings EMG linear envelope (Figure 5) nor the hamstrings contribution to KCF (Figure 6b) increased during late stance. The only significant difference in lower-extremity sagittal joint kinematics or kinetics was a 0.72±0.63%BW*height (31±24%, p=0.015) reduction in the late-stance knee flexion moment (Figure 7). This finding aligns with our observed reduction in gastrocnemius activation but unchanged hamstrings activation.

**Figure 6:**
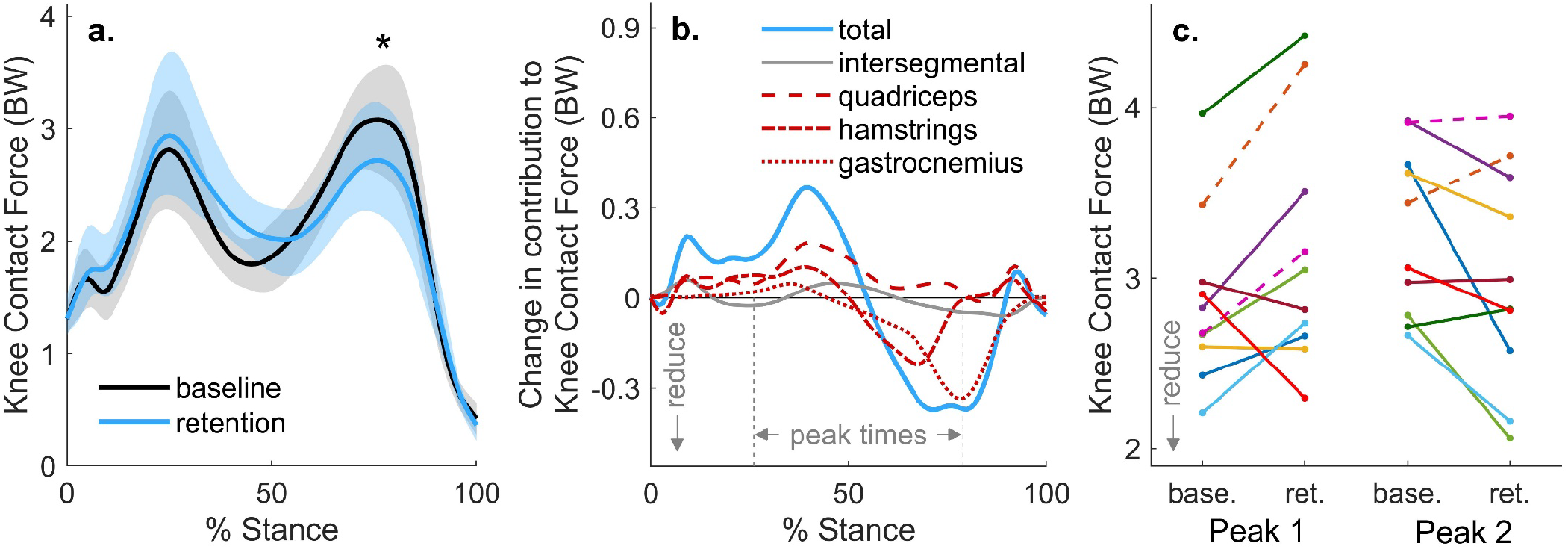
**(a)** The average (line) and standard deviation (shading) of knee contact force for the participants (n=8) who reduced late-stance gastrocnemius activation during the retention trial. These participants reduced their second peak of knee contact force by 0.38±0.39 times bodyweight (BW), or 12±12% (p=0.029) during the retention trial, compared to baseline. **(b)** For the same eight subjects, the change in knee contact force is broken down into the intersegmental and muscle force components. A 0.25±0.30 BW reduction in gastrocnemius force at the time of second peak knee contact force was the main contributor to the reduction in second peak contact force. **(c)** Changes in the first and second peak contact force between baseline (base.) and retention (ret.) are shown for all participants (n=10), with the two participants who did not retain a reduction in late stance gastrocnemius activity represented with dashed lines. Six of the eight individuals who reduced late-stance gastrocnemius activation reduced their second peak knee contact force, but five increased their first peak.

**Figure 7:**
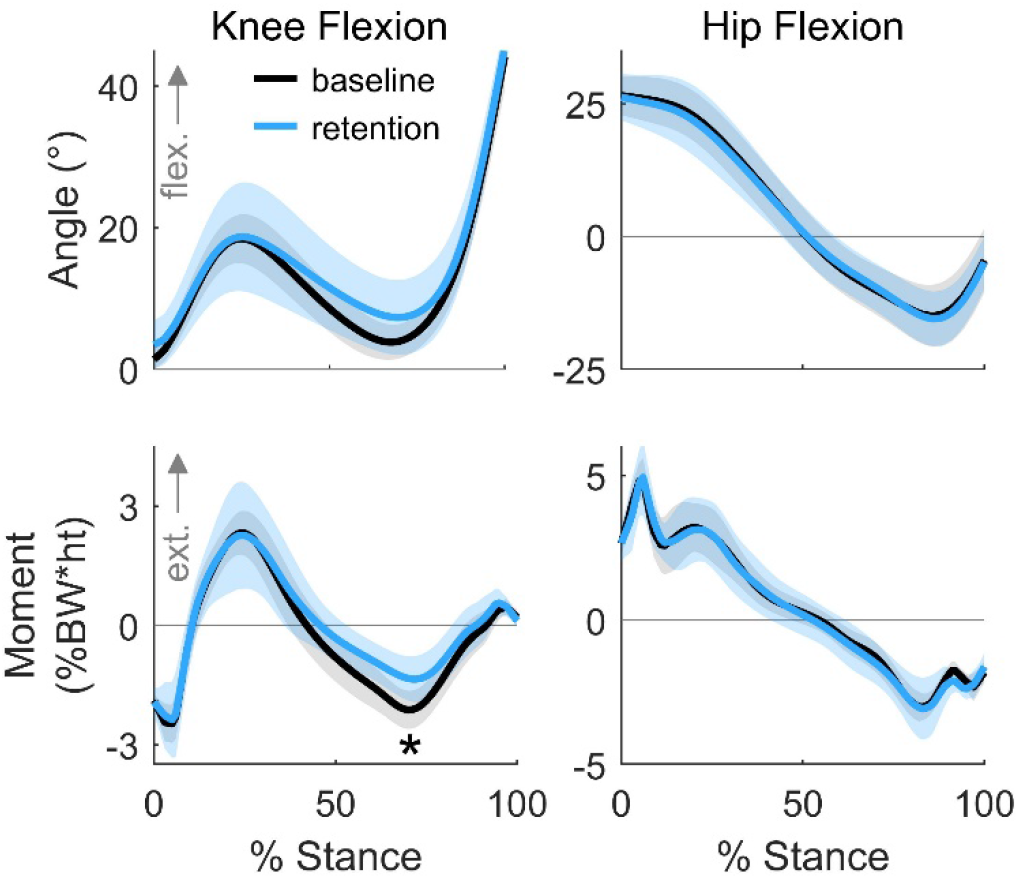
The average (line) and standard deviation (shading) of knee and hip kinematics and kinetics for the participants (n=8) who retained a reduction in their late-stance gastrocnemius activation during the retention trial. During the retention trial, participants walked with a 0.72±0.63%BW*ht (31±24%, p=0.015) smaller late-stance knee flexion moment. There was large variance in the first peak knee angle and moment during the retention trial. Peak hip kinematics and kinetics were not significantly different during early or late stance.

## Discussion

We developed a muscle coordination retraining intervention that reduces knee loading. We first used musculoskeletal simulations to identify simple changes in coordination that reduce knee contact force, which informed the design of a biofeedback intervention. With simple biofeedback of their muscle activity, healthy individuals were able to quickly change the relative activation of their gastrocnemius and soleus muscles while still generating a similar ankle plantarflexion moment. This change in coordination resulted in a 12% reduction in simulation-estimated knee contact force that may have therapeutic benefit for individuals with knee osteoarthritis. These results suggest that simulation-based intervention design is a promising tool for identifying new, learnable coordination strategies that achieve a therapeutic objective.

Our results build upon prior EMG biofeedback work by demonstrating that humans can quickly alter the coordination of redundant muscles during a complex task with simple feedback. Acute coordination changes have been learned within a single visit for static activities^28^, but clinical coordination retraining programs can last many weeks^31^. Our participants reduced their gastrocnemius-to-soleus activation ratio during walking after only five minutes of feedback. This suggests that multiple coordination strategies could be taught during a single visit for applications, like human-in-the-loop optimization, in which users need to rapidly adapt to new conditions^38^. Prior work has also demonstrated that precise control of individual motor units or muscles is more easily attained during isometric contractions^28,30^ or controlled physical therapy activities^31,32^, but this study suggests that muscle coordination can be retrained during dynamic activities like walking. Further studies are necessary to determine how complex of coordination changes can be taught during functional activities.

Pizzolato et al. showed that during a single visit, healthy individuals were not able to identify coordination patterns that reduced KCF with real-time feedback of KCF alone^39^. This could be due to the complexity of the dynamics that relate muscle coordination to KCF. When given feedback of the activation of two muscles, most of our participants reduced KCF, which highlights the importance of selecting biofeedback targets that are both simple and effective.

The ability of a gastrocnemius avoidance coordination pattern to reduce knee contact force suggests that coordination retraining could be used to reduces joint contact forces other arthritic joints throughout the body. The 0.38BW or 12% reduction in KCF_P2_ is similar to the joint-offloading effect of losing 15-38% of bodyweight^40–42^. The reduction in KCF_P2_ is similar to what has been achieved with kinematic gait modifications^43,44^ or assistive devices^14,43,45^, but does not require conspicuous kinematic or lifestyle changes that often hinder patient adoption^17,46^. By leveraging functional differences between redundant muscles throughout the musculoskeletal system, coordination retraining is a flexible way to reduce loading. It could be used to shift loading between the compartments of the knee or to reduce loading in other joints, like the hip^24^, for which there are a limited number of other offloading interventions.

In addition to designing interventions to reduce joint contact force, musculoskeletal simulations can identify coordination retraining interventions that optimize for other important clinical and sports-performance metrics. Simulations can identify optimal muscle coordination strategies for walking or running with a robotic assistive device^47,48^. Providing simulation-guided EMG biofeedback may augment the natural motor learning process, allowing a device user to more quickly converge on an energetically-favorable coordination strategy. Additionally, muscle coordination patterns identified by simulations could improve sports performance. For example, coordination patterns that maximize the distance of a long jump^49^ could be taught to athletes with simplified EMG biofeedback. Finally, this framework could become an important injury prevention and rehabilitation tool. For example, new coordination patterns could be designed to help reduce strain in the anterior cruciate ligament when cutting or in the ulnar collateral ligament during baseball pitching. Even after an injury, new muscle coordination patterns may help normalize joint mechanics^23^, potentially making coordination retraining an important part of post-injury rehabilitation.

It is important to acknowledge the limitations of this study. We demonstrate that healthy individuals can learn and retain a new coordination pattern in the context of a single visit, but further work is necessary to evaluate learning and retention over long durations in free-living conditions. Although there are challenges with day-to-day EMG normalization, EMG-based biofeedback is amenable to a wearable solution for in-home training. Second, we show the feasibility of retraining muscle coordination in young, healthy participants. Individuals with osteoarthritis may find this modification more challenging to learn due to impairments in muscle strength or motor control. Third, on average, our participants did not maintain their natural walking kinetics when adopting the gastrocnemius avoidance coordination pattern. Our feedback successfully taught individuals to change ankle plantarflexor muscle activity while maintaining a natural ankle plantarflexion moment; however, we did not give feedback to increase hamstrings activation, resulting in a reduction in the late-stance knee flexion moment. If future studies aim to retain identical joint kinematics and kinetics, additional feedback should be given to instruct participants to achieve all of the necessary compensatory changes in muscle activation. Fourth, although six of the eight subjects who reduced their late-stance gastrocnemius activity reduced the second peak of KCF, five of these individuals increased their first peak of KCF. Future studies that investigate this gait pattern should monitor changes in kinetics and muscle activity during early stance that affect the first peak of KCF. Finally, we estimated changes in KCF using a musculoskeletal model instead of directly measuring them with an instrumented knee implant. However, a similar static optimization approach has been shown to accurately detect gait modification-induced changes in KCF measured with an instrumented knee implant^50^. The changes in simulated muscle activation between natural and gastrocnemius avoidance gaits also matched changes in EMG (Figure 5), providing confidence in the accuracy of our simulated changes in KCF.

In summary, humans can learn and retain changes to their muscle coordination pattern during walking when given simple biofeedback. We taught healthy adults to alter the activation of their redundant plantarflexor muscles, which reduced knee loading, demonstrating that coordination retraining may be a valuable new conservative treatment to offload osteoarthritic joints. More generally, simulation-inspired muscle coordination retraining may be an effective way to teach people novel, therapeutic coordination strategies. This approach capitalizes on the adaptability of the human neuromusculoskeletal system to optimize muscle coordination, with compelling injury prevention, rehabilitation, and sports performance applications.

## Methods

### Simulation-based intervention design

Previous work had suggested that knee contact force could be reduced with a gastrocnemius avoidance coordination pattern^20^, but it was unclear what compensatory changes in muscle activation were necessary. To address this question, we simulated one gait cycle of normal walking kinematics (healthy male, speed 1.25m/s) with two feasible muscle coordination patterns: a natural pattern that minimized a surrogate for energy expenditure and a gastrocnemius avoidance pattern that minimally activated the gastrocnemius (Figure 2a). We used OpenSim 4.0^51,52^ and a custom static optimization implementation in MATLAB R2017b (Mathworks, Inc., Natick MA, USA) for musculoskeletal modeling and simulation.

We modeled the lower extremities and torso using the musculoskeletal model described by Rajagopal et al.^35^ with 21 degrees of freedom, 80 musculotendon actuators for the lower extremities, and three ideal torque actuators for the torso. The model included six degrees of freedom between the pelvis and the ground, three rotational degrees of freedom between the pelvis and torso, three rotational degrees of freedom at the hip, one rotational degree of freedom at the knee that parametrized the remaining rotational and translational degrees of freedom of the tibiofemoral^53^ and patellofemoral^54^ joints, and one rotational degree of freedom at each of the ankle and subtalar joints. We modified the model by calibrating the each muscle’s passive muscle force-length curves so that joint moments generated by passive muscle forces more closely matched experimental data^55^. We also altered the muscle paths of the hip abductor musculature to more closely match moment arms estimated from experiments^56,57^, finite element models^58^, and MRI^59^ (See Supplemental Material). The modified model was scaled to match the anthropometric measurements taken from the static trial, and virtual markers on the model were moved to match experimental marker locations during this trial. Model kinematics were estimated using the Inverse Kinematics tool in OpenSim, which minimizes the error between the positions of experimental markers and virtual model markers. Joint moments were computed using the Inverse Dynamics tool in OpenSim with low-pass-filtered kinematics (6Hz, 6^th^ order, zero-phase shift Butterworth) and ground reaction forces (6Hz, 4^th^ order, zero-phase shift Butterworth) as inputs.

We developed a custom static optimization algorithm in MATLAB, using the OpenSim API, to solve the muscle redundancy problem. Two objective functions were used. The first minimized the sum of squared muscle activations (Equation 1), which is a commonly-used surrogate for metabolic cost^33,34^. The second was a gastrocnemius avoidance objective function with a heavily-weighted penalty on gastrocnemius activation^20^ (Equation 2). Muscle-generated joint moments were constrained to match joint moments from inverse dynamics (Equation 4) at each timestep (*t_k_*).

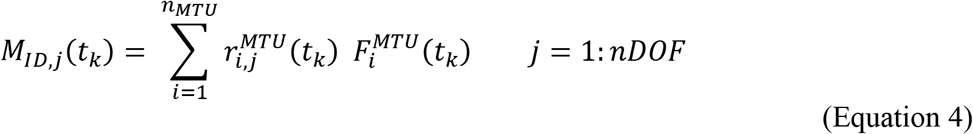

where *M_ID,j_* are the inverse dynamics joint moments for each of *j* degrees of freedom (DOF), 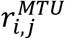 are the moment arms^60^ of the *i^th^* musculotendon actuator (MTU) about the *j^th^* DOF, and 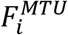 are the MTU forces. MTU forces were computed using the Hill-type model^61^ described by Millard et al.^62^. We approximated tendon compliance by solving the musculotendon static equilibrium equation (Equation 5) for muscle fiber length using the MTU length from the current step and the activation from the previous step:

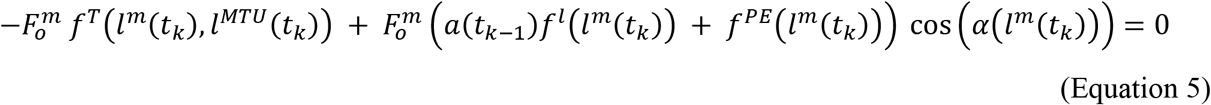

where 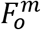 is the optimal muscle force, *f^T^*(*l^M^, l^MTU^*) is the tendon force-length multiplier as a function of the muscle fiber and MTU lengths, *α* is muscle activation between 0 and 1, *f^l^* (*l^m^*) is the active muscle fiber force-length multiplier, *f^PE^*(*l^m^*) is the passive muscle fiber force-length multiplier, and *α*(*l^m^*) is the muscle pennation angle. After solving Equation 5 for *l^m^*(*t_k_*) using Newton’s method^62^, we fixed *l^m^*(*t_k_*) and solved for *F^MTU^*(*t_k_*) as a function of the design variable, *α*(*t_k_*), for each muscle (Equation 6).

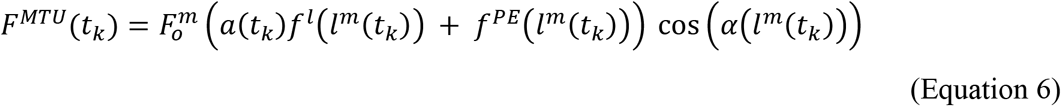

We then used *fmincon* in MATLAB to solve the static optimization problem with the combination of Equations 4 and 6 as a constraint and either Equation 1 or Equation 2 as the objective function. The static equilibrium computation in Equation 5 does not account for the force-velocity property of muscle. However, in pilot testing, we found that static optimization solutions that used a compliant tendon but excluded the force-velocity property of muscle yielded activations that more closely matched EMG than solutions that assumed a rigid tendon but incorporated the force-velocity property. This improvement was especially evident in muscles, like the gastrocnemius and soleus, that have tendons that are several times longer than their optimal muscle fiber lengths.

Using the muscle forces from static optimization, we used the Joint Reaction Analysis tool in OpenSim to compute KCF as the reaction force along the longitudinal axis of the tibia. We also compared the differences in muscle forces in the major muscle groups that generate the late-stance ankle plantarflexion, knee flexion, and hip flexion moments: the soleus, the gastrocnemius (medial and lateral heads), the hamstrings (semitendinosus, semimembranosus, biceps femoris long and short head), the iliopsoas (iliacus, psoas), and the rectus femoris. Finally, after evaluating the changes in muscle force between coordination patterns, we chose to focus biofeedback on the two muscle groups with the greatest changes in force: the gastrocnemius and soleus. By reducing gastrocnemius activation and increasing soleus activation, an individual could, in theory, reduce the gastrocnemius contribution to knee contact force without changing the net ankle plantarflexion moment.

### Muscle coordination retraining experiment

Eleven individuals enrolled in the study and 10 (4 female, 26±4 years old, 22.8±2.1 BMI) completed it after providing informed consent to a protocol approved by the Stanford University Institutional Review Board. We included individuals who did not have a history of lower-limb injury in the past year. Before individuals began the walking portion of the study, we evaluated whether our surface EMG electrodes could measure different signals between the medial gastrocnemius and soleus. We asked participants to perform standing and seated plantarflexion exercises, since these activities induce large changes in the relative activation of the gastrocnemius and soleus muscles due to varying amounts of knee flexion^63^. Because our target reduction in gastrocnemius-to-soleus activation ratio during walking was 15%, we only included participants who reduced their activation ratio by at least 15% during the seated plantarflexion activity compared to the standing plantarflexion activity. One participant was excluded for this reason.

Participants completed a single visit to a motion capture laboratory with a force-instrumented treadmill (Bertec Corporation, Columbus OH, USA), an 11-camera motion capture system (Motion Analysis Corporation, Santa Rosa, CA, USA), and a wireless surface EMG system (Delsys Corp., Natick, MA, USA). We placed markers bilaterally on the 2nd and 5th metatarsal heads, calcanei, medial and lateral malleoli, medial and lateral femoral epicondyles, anterior and posterior superior iliac spines, acromion processes, sternoclavicular joints, and on the C7 vertebrae. Markers on the medial femoral epicondyles and malleoli were removed prior to walking trials, and 16 additional markers were used to aid in limb tracking. After randomly selecting a leg for analysis and biofeedback, we placed nine EMG electrodes unilaterally on the soleus (medial aspect), medial gastrocnemius, lateral gastrocnemius, tibialis anterior, biceps femoris, semitendinosus, rectus femoris, vastus medialis, and vastus lateralis.

Prior to walking, participants performed maximum voluntary contraction activities for EMG normalization, static and dynamic calibration trials to aid in scaling of the musculoskeletal model, and calf-raises to evaluate EMG sensor placement. After walking on the treadmill to warm up, participants performed two resisted isometric and isokinetic maximum voluntary contraction activities: prone knee flexion and supine ankle dorsiflexion. Participants then performed five maximum effort hip flexion kicks and five maximum height jumps^64^. We calculated EMG linear envelopes by bandpass filtering (30-500Hz, 4^th^ order, zero-phase shift Butterworth), rectifying, and low-pass filtering (6Hz, 4^th^ order, zero-phase shift Butterworth) the raw EMG signals. We normalized all future EMG linear envelopes by the maximum value for each muscle measured during any of the maximum voluntary contraction activities. Participants then performed a standing static calibration trial for model scaling and hip circumduction trials for the identification of functional hip-joint-center locations^65^. To familiarize participants with the anatomy of their triceps surae muscles, we showed them images of the gastrocnemius and soleus muscles and palpated both muscles. To allow participants to feel the difference between gastrocnemius and soleus activation and to evaluate the differences in EMG signals, participants performed two sets of 10 double-leg calf raises. The participants were standing (hip and knee at 0°) during the first set to primarily activate the gastrocnemius, and were seated (hip and knee at 90°) with 4.5 kg weights on each knee during the second set to primarily activate the soleus. The average reduction in gastrocnemius-to-soleus activation ratio during the seated trial, compared to the standing trial, was 68%±14% for the 10 individuals who completed the study and 11% for the individual who was excluded from the study for the inability to reduce the activation ratio by at least 15% during the seated plantarflexion.

After acclimating to walking on the treadmill at 1.25ms^−1^ for five minutes, participants performed five six-minute walking trials: a baseline trial, three feedback trials, and a retention trial (Figure 2b). During the baseline walking trial, they were instructed to walk naturally. We computed their baseline gastrocnemius-to-soleus activation ratio (Equation 3) and baseline stance-phase-averaged gastrocnemius EMG linear envelope. We averaged these values over the final minute of the baseline trial and normalized real-time measurements by the average baseline values in subsequent trials. During the first feedback trial, we used a real-time bar plot to instruct participants to reduce their activation ratio by at least 15% relative to baseline while maintaining normal walking kinematics. We instructed them to explore different strategies during the first four minutes of the trial, to converge on their most successful strategy during the fifth minute, and to maintain this strategy after feedback was removed for the sixth minute. During the second and third feedback trials, participants were instructed to continue reducing their activation ratio below the target line, which was set to either a 15% reduction from baseline or their average reduction from baseline during the previous trial, whichever was greater. The 15% target was chosen as a feasible goal from pilot testing, but the adaptive target-setting method allowed for increasingly challenging goals for participants who were successful during the previous trial. During these final two trials, participants were given additional feedback of their average gastrocnemius activity, represented by the color of the bar. We instructed them to reduce their activation ratio by reducing gastrocnemius activation, or to achieve a small, blue bar (Figure 2b). During the retention trial, we removed the feedback but instructed participants to walk with the coordination pattern that they learned during the feedback trials. We analyzed the final 30 steps of the sixth minute of all trials. For analysis, we used the same EMG filtering and normalization processes as used for the real-time experiment, and we averaged muscle activity over the duration of the stance phase.

### EMG-informed simulation of knee contact force

We simulated these experimental data to estimate joint kinematics, kinetics, and KCF using OpenSim^52^ and the previously-described custom static optimization implementation. For simulations, five gait cycles were selected from each of the baseline and retention trials that had the smallest absolute differences in activation ratio compared to the average value from the final 30 steps in the trial.

Changes in plantarflexor EMG were incorporated into the static optimization simulation by constraining the simulated gastrocnemius-to-soleus activation ratio to match the ratio measured with EMG. First, a 40ms electromechanical delay^66^ was added to the medial gastrocnemius, lateral gastrocnemius, and soleus EMG linear envelopes. The simulated medial gastrocnemius-to-soleus activation ratio was constrained to match the EMG ratio within 2% (Equation 7). A similar constraint was enforced for the lateral gastrocnemius-to-soleus activation ratio.

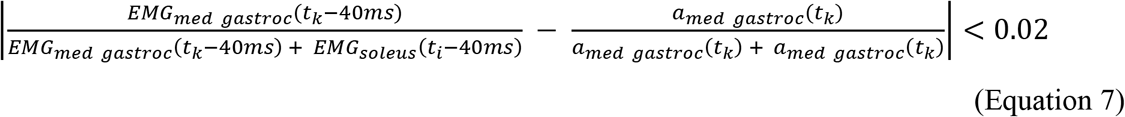

To validate our simulations, we qualitatively compared simulated activations to EMG linear envelopes with the 40ms electromechanical delay (Figure 5). For this comparison, we computed weighted averages of simulated activation and EMG linear envelopes for the vasti (vastus medialis and vastus lateralis) and biarticular hamstrings (semitendinosus and biceps femoris long head). The optimal muscle force from the musculoskeletal model was used as the weight for each muscle.

We used the static optimization solutions and the Joint Reaction Analysis tool to compute KCF and decompose it into its muscle and intersegmental force components. We found the intersegmental reaction force by computing KCF from a simulation with all degrees of freedom actuated by ideal torque actuators. To elucidate the contribution of different muscle groups to KCF (Figure 1, Figure 6b), we prescribed the static optimization-based muscle forces to a subset of muscles, generated the remaining net joint moments with ideal torque actuators, and subtracted the intersegmental force from the computed KCF. We defined the following functional groups of knee-crossing muscles: the quadriceps (vastus medialis, vastus intermedius, vastus lateralis, and rectus femoris), the hamstrings (biceps femoris long and short heads, semitendinosus, semimembranosus, gracilis, and sartorius), the gastrocnemius (medial and lateral gastrocnemii), and the tensor fascia latae.

### Statistics

All statistical analyses were performed in MATLAB (R2017b) unless otherwise noted. After testing data for normality using the Shapiro Wilk test^67^, we compared normally distributed data using a paired sample, two-sided t-test and compared non-normally distributed data using a two-sided Wilcoxon signed rank test. For comparisons of activation ratio and muscle activation changes across the feedback and retention trials, we report p-values after controlling for the false detection using R (v3.5.3, R Foundation for Statistical Computing, Vienna, Austria)^68,69^. We used a two-one-sided test for equivalence using the tool provided by Lakens et al.^70^ with equivalence bounds of one baseline standard deviation to compare the average ankle plantarflexion moment between the baseline and retention trials. We defined peaks in the knee and hip joint angle and moment curves as the maximum value during the first 50% of the stance phase and the minimum value during the final 50% of the stance phase. Due to the exploratory nature of evaluating changes peak angles and moments between baseline and retention, we did not correct for multiple comparisons. Values reported are mean ± standard deviation, and α=0.05.

## Data Availability

The experimental data and simulation results are available at https://simtk.org/projects/coordretraining. The modified musculoskeletal model is available at https://simtk.org/projects/fbmodpassivecal.

## Code Availability

The static optimization code and code for calibrating the passive muscle force curves in a musculoskeletal model are available at https://github.com/stanfordnmbl/MatlabStaticOptimization and https://github.com/stanfordnmbl/PassiveMuscleForceCalibration, respectively. The real-time EMG biofeedback code is specific to our motion capture system, and will be shared upon reasonable request to the authors.

## Author Contributions

SDU and SLD conceived of the study. SDU performed the musculoskeletal simulations, collected the experimental data, and analyzed the results. AS advised with the development of simulation software. All authors aided in data interpretation. SDU drafted the manuscript, and all authors critically revised and approved the final manuscript.

## Acknowledgements

The authors would like to thank Nick Bianco, Carmichael Ong, and Amy Silder for their advice on the simulations and experiments. SDU was funded by a fellowship from the National Science Foundation (DGE-114747) and the Sang Samuel Wang Stanford Graduate Fellowship from the Stanford Office of the Vice Provost for Graduate Education. This work was also supported by grant P41 EB027060 from the National Institutes of Health.

## Competing Interests

Stanford University has applied for a patent on behalf of SDU and SLD describing the muscle coordination retraining technique, entitled *Real-time electromyography feedback to change muscle activity during complex movements*. The patent is pending at the time of manuscript submission. The authors have no other competing interests to disclose.

## Notes

https://simtk.org/projects/coordretraining

https://simtk.org/projects/fbmodpassivecal

https://github.com/stanfordnmbl/MatlabStaticOptimization

https://github.com/stanfordnmbl/PassiveMuscleForceCalibration

