## Supplemental Materials for "Muscle coordination retraining inspired by musculoskeletal simulations: a study on reducing knee loading"

**Supplemental Material**

*Hip abductor muscle path adjustments*

We adjusted the origin and insertion points of the hip abductors in the model described by Rajagopal et al.[35] to improve estimates of hip flexor muscle activation (Figure S1). During simulations of normal walking, static optimization was requiring non-physiologically large iliacus and psoas activations despite hip moments that matched normative values in the literature. The moment-generating capacity of the hip flexors was validated by Rajagopal et al.; however, this study investigated the sagittal plane moment in isolation, not in combination with frontal or transverse plane moments. We determined that the high hip flexor muscle activity may have, in part, resulted from the combined hip flexion and abduction moments that are generated during the latter half of stance phase of walking. When this model is in hip extension, all six of the gluteus medius and gluteus minimus muscle fibers, which generate the majority of the hip abduction moment, have sagittal plane moment arms that extend the hip (Figure S2). During late stance, these muscles generate an antagonistic hip extension moment that requires elevated activation of the iliacus and psoas. Experimental studies[56,57], finite element models[58], and other MRI-based musculoskeletal models[59] suggest that the anterior fibers of the gluteus medius and minimus generate a hip flexion moment when the hip is in an extended position. We adjusted the origin and insertion points of the gluteus medius, gluteus minimus, and tensor fascia latae to more closely match these data and models (Figure S2).

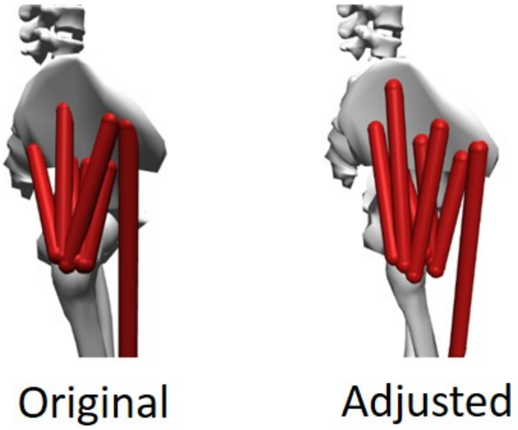


**Figure S1**: Changes in the origin and insertion points of the hip abductors from the original Rajagopal et al.[35] model (left) to the adjusted model (right). The origin of the gluteus medius and minimus were moved superiorly and laterally to increase the abduction moment arm to match experimental data. The insertion of the gluteus medius and minimus were moved anteriorly to allow them to generate a greater flexion moment to match experimental and model-based moment arms (Figure S2).

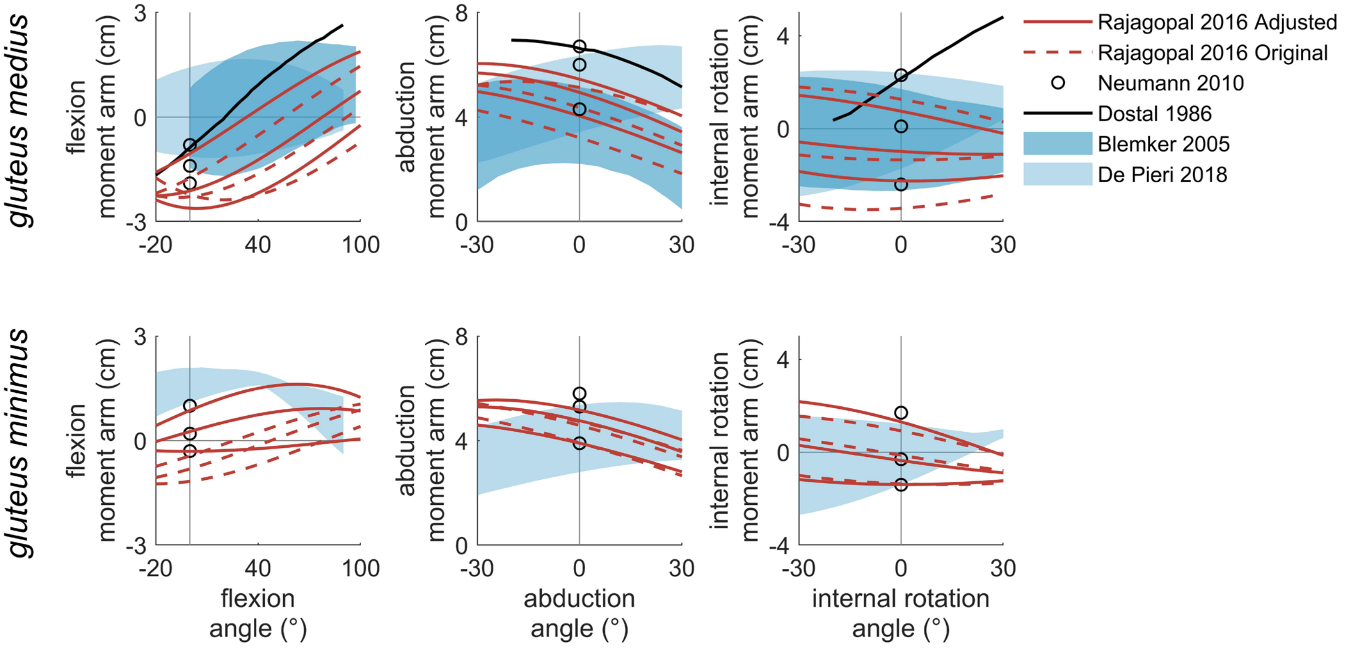


**Figure S2**: The original and adjusted moment arms for the musculoskeletal model described in Rajagopal et al.[35] compared to moment arms from experiments (Dostal et al.[56], Neumann[57]), finite element models (Blemker et al.[58]), and MRI (De Pieri et al.[59]). The Dostal et al. moment arm curve is from the anterior region of the gluteus medius, and all other moment arms are from a range of fibers from the posterior to the anterior regions of the muscles.

*Passive muscle force calibration*

The muscles in the Rajagopal et al.[35] model generate passive joint moments that are larger than those that have been measured in experiments[55], especially at large knee or hip flexion angles[71] (Figure S3). Previous studies have addressed this by changing the muscle optimal fiber lengths, tendon slack lengths, or muscle geometry[71]. These changes were driven by passive muscle forces but they also affect the active force-generating capacity of the muscle. Another approach to modifying the passive muscle forces is to modify the passive muscle force-length curve, which does not affect the active force-generating capacity of the muscle.

We calibrated the passive force-length curves for each muscle in the Rajagopal model to more closely match experimentally-measured passive joint moment curves[55]. Using constrained optimization in MATLAB, we minimized the root mean square difference between passive joint moment curves from the model and experiment. As design variables, we used two of the values that parameterize the passive force-length curve in the Millard et al.[62] muscle model: the muscle length ($l^m$) when it begins to generate force (default value: $l_o^m$) and the length at which passive muscle force reaches its optimal fiber force ($F_o^m$, default value: 1.7 $l_o^m$). We limited changes in $l^m$ values to 0.2 $l_o^m$ above or below their default values. The resulting changes in passive joint moment curves are shown in Figure S3 and in passive force-length curve parameters in Figure S4 and Table S1. After calibration, the quadriceps and gluteus maximus muscles all began generating passive muscle forces at longer muscle fiber lengths, which reduces the passive moments generated at large knee and hip flexion moments. The iliacus and psoas began generating passive force at shorter muscle fiber lengths, which allows them to generate a larger passive hip flexion moment when the hip is extended. Incorporating calibrated passive muscle forces into our static optimization tool helped reduce large hip flexor muscle activations during stance phase as well as quadriceps-hamstrings co-contraction when the knee is flexed during swing phase.

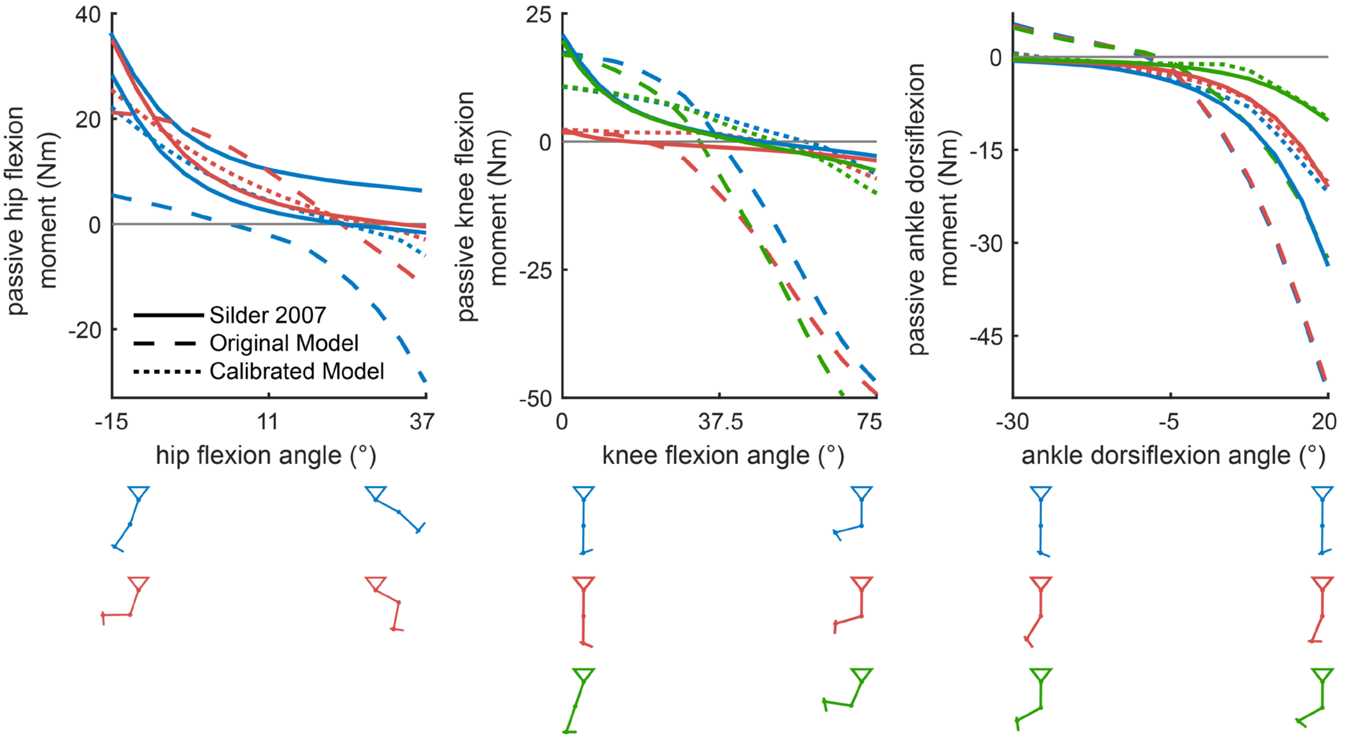


**Figure S3**: Sagittal plane passive joint moment curves for the hip, knee, and ankle. Calibration improved the agreement in passive joint moments between the Rajagopal et al.[35] model and experimentally-measured moments from Silder et al.[55]. Each joint was moved over the shown range of motion with other joints fixed at various angles (see Silder et al.). For example, the hip was moved from 15° of extension to 37° of flexion (left) with the knee fixed at 15° (blue) and 60° (red).

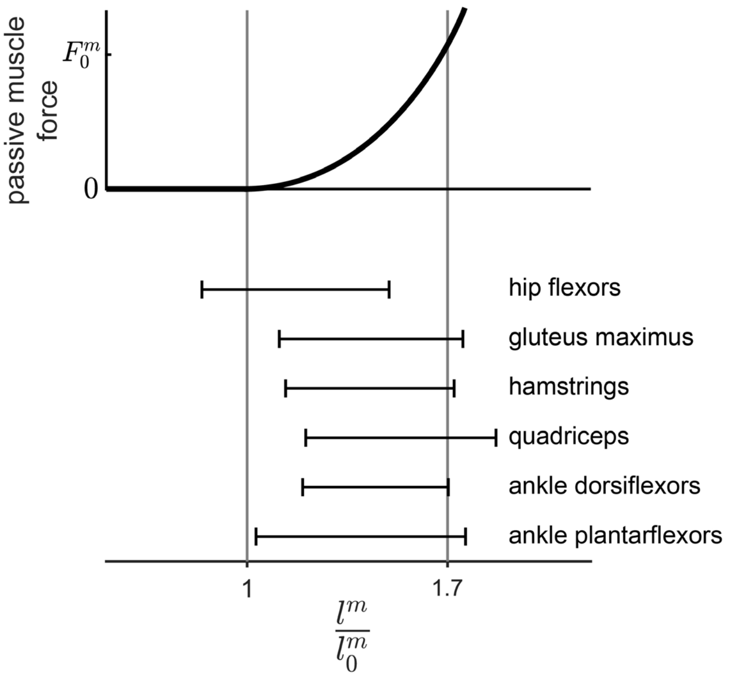


**Figure S4**: Calibrated normalized muscle fiber lengths that parameterize the passive muscle force curve for sagittal plane muscle groups. At $F^m = 0$, the default $l^m/l_o^m = 1$. At $F^m = F_o^m$, the default $l^m/l_o^m = 1.7$. Muscle groupings are defined in Table S1. With the exception of the hip flexors, most muscle groups begin generating passive force at a longer fiber length after calibration.

**Table S1**: Calibrated muscle fiber lengths that parameterize the passive muscle force curve for each muscle in the Rajagopal et al.[35] model.

| | $\frac{l^m}{l_o^m}$ at $F^m = 0$ | $\frac{l^m}{l_o^m}$ at $F^m = F_o^m$ |
|---|---|---|
| **default value** | 1 | 1.70 |
| hip flexors | | |
| psoas | 0.93 | 1.50 |
| iliacus | 0.88 | 1.50 |
| addlong | 0.82 | 1.50 |
| tfl | 1.20 | 1.72 |
| gluteus maximus | | |
| glmax1 | 0.92 | 1.84 |
| glmax2 | 1.20 | 1.72 |
| glmax3 | 1.20 | 1.71 |
| hamstrings | | |
| bflh | 1.20 | 1.71 |
| bfsh | 1.14 | 1.70 |
| semimem | 1.20 | 1.75 |
| semiten | 1.18 | 1.71 |
| grac | 0.80 | 1.66 |
| sart | 0.80 | 1.50 |
| quadriceps | | |
| vasmed | 1.20 | 1.90 |
| vaslat | 1.20 | 1.90 |
| vasint | 1.20 | 1.90 |
| recfem | 1.20 | 1.90 |
| ankle dorsiflexors | | |
| tibant | 1.19 | 1.71 |
| ankle plantarflexors | | |
| gasmed | 1.20 | 1.90 |
| gaslat | 1.20 | 1.67 |
| soleus | 1.20 | 1.90 |
| tibpost | 0.80 | 1.50 |
| other | | |
| addbrev | 0.96 | 1.67 |
| addmagDist | 1.00 | 1.70 |
| addmagIsch | 1.01 | 1.70 |
| addmagMid | 1.00 | 1.70 |
| addmagProx | 1.00 | 1.70 |
| glmed1 | 1.00 | 1.70 |
| glmed2 | 1.06 | 1.70 |
| glmed3 | 1.12 | 1.70 |
| glmin1 | 1.00 | 1.70 |
| glmin2 | 1.00 | 1.70 |
| glmin3 | 1.01 | 1.70 |
| piri | 1.11 | 1.70 |
| perbrev | 0.92 | 1.69 |
| perlong | 0.80 | 1.50 |
| edl | 1.12 | 1.72 |
| ehl | 1.09 | 1.73 |
| fdl | 0.95 | 1.70 |
| fhl | 1.04 | 1.70 |
